# *Pelagibacter* phage Skadi - An abundant polar specialist that exemplifies ecotypic niche specificity among the most abundant viruses on Earth

**DOI:** 10.1101/2022.08.10.503363

**Authors:** Holger H. Buchholz, Luis M. Bolaños, Ashley G. Bell, Michelle L. Michelsen, Michael J. Allen, Ben Temperton

**Affiliations:** School of Biosciences, University of Exeter, Exeter, UK

## Abstract

Bacteria in the SAR11 clade are the most abundant members of surface marine bacterioplankton and are a critical component of global biogeochemical cycles. Similarly, pelagiphages that predate SAR11 are both ubiquitous and highly abundant in the oceans. These viruses are predicted to shape SAR11 community structures and increase carbon turnover throughout the oceans. Yet, ecological drivers of host and niche specificity of pelagiphage populations are poorly understood. Here we report the global distribution of a novel pelagiphage called Skadi isolated from the Western English Channel using a cold-water ecotype of SAR11 (HTCC1062) as bait. Skadi is closely related to the globally dominant pelagiphage HTVC010P. We show that, along with other HTVC010P-type viruses, Skadi belongs to a distinct viral family within the order *Caudovirales* for which we propose the name *Ubiqueviridae*. Metagenomic read recruitment identifies Skadi as one of the most abundant pelagiphages on Earth. *Skadi* is a polar specialist, replacing HTVC010P at high latitudes. Experimental evaluation of Skadi host-range against cold- and warm-water SAR11 ecotypes supported cold-water specialism. Read mapping from the Global Ocean Viromes project (GOV2) showed that relative abundance of Skadi correlated negatively with temperature, and positively with nutrients, available oxygen and chlorophyll concentrations. In contrast, relative abundance of HTVC010P correlated negatively with oxygen and positively with salinity, with no significant correlation to temperature. The majority of other pelagiphages were scarce in most marine provinces, with a few representatives constrained to discrete ecological niches. Our results suggest that pelagiphage populations persist within a global viral seed-bank, with environmental parameters and host availability selecting for a few ecotypes that dominate ocean viromes.

## Introduction

Bacteriophages are the most abundant organisms in the oceans and play a critical role in driving microbial global biogeochemical cycles, particularly carbon cycling [1–4]. During an infection, phages hijack their hosts, alter cellular metabolism and shape the structure of microbial populations [5–7]. Ultimately, phages predate and kill bacterial host cells, releasing the carbon and other cell-bound nutrients back into the environment, thereby recycling organic compounds [3, 8]. These compounds are remineralised by other microbes within the ‘*viral shunt*’, which is thought to be an essential part of the marine food web [3, 9]. Phages drive microbial evolution and are agents of frequent genetic exchange between hosts as well as other phages, creating highly diverse genomic populations [10, 11]. In marine systems, discrete viral populations persist within spatial and temporal niche, in which the majority of sequence variation occurs in metabolic genes, indicating niche specificity [12, 13]. Genomic clusters for T4-like phages infecting Cyanobacteria, were found to exhibit spatial and temporal abundance patterns suggesting that they represent distinct viral ecotypes [13]. Viral communities at the Bermuda Atlantic Time-Series (BATS) station also revealed recurrent seasonal patterns linking viral communities with stratification and mixing events in the water column [14]. Temporal analysis of T4-like phages and their hosts at the San Pedro Ocean Time series station further suggested that the relative abundance of virus populations is tightly linked to those of their associated hosts [15]. Therefore, spatiotemporal distribution and ecotypes of viral populations are interlinked and influenced by both the physical and biological parameters that shape the ecotypic specificity of their hosts [16].

The ubiquitous alphaproteobacterial order *Pelagibacterales* (clade SAR11) comprise up to one-third of total marine microbial communities, making them likely the most abundant bacteria on Earth [17, 18]. SAR11 are an ancient and diverse clade, characterised by small, streamlined genomes with low G+C % content, high gene synteny and are phylogenetically organised into specialized temperature-associated ecotypes [19–21]. Phages infecting members of the SAR11 (pelagiphages) are the most abundant viruses in global ocean viromes [22, 23] and have been isolated across most major oceans, resulting in 40 different cultivated species, including 37 of podophage morphotypes [22–27], two myophages [22, 25] and one siphophage [26]. Owing to the challenges of culturing SAR11 [28, 29] and isolating phages for them [26], experimental evaluation of predicted pelagiphage ecology is in its infancy. Read recruitment from global viral metagenomes to genomes of pelagiphages has identified several pelagiphages with abundant and cosmopolitan distributions irrespective of temperature (such as HTVC010P [22], HTVC023P [23], and vSAG-37-F6 [30, 31]). However, when conservative read recruitment standards are applied, including a minimum genome coverage cutoff to avoid false discovery rate of 20-25% [32], the vast majority of pelagiphages (22 out of 32 with mean RPKM <100, 31 out of 32 with median RPKM of zero [26]) have low abundance distributed across few geographic locations, in support of the viral seed-bank hypothesis [33], which states that ocean currents distribute phages across the globe, but only a fraction of phages in the ocean are actively replicating, depending on local host availability. HTVC010P remains the sole representative of cultured pelagiphages with high abundance in epipelagic samples from both temperate tropical and polar regions.

Two possibilities may explain the extraordinary abundance of HTVC010P across the global oceans: One hypothesis is that HTVC010P has a broad host range and is capable of infecting multiple host ecotypes and are thus propagated in both warm and cold-water environments. Experimentally confirmed host range experiments for HTVC010P do not exist, host range data for the majority of other pelagiphages is limited [34], but existing evidence suggests that they tend to have narrow host ranges restricted to specific ecotypes [23, 24, 26]), at least under laboratory conditions. To date, only a single, low abundance pelagiphage (*Pelagibacter* phage Bylgja) has been found to experimentally infect both warm and cold-water ecotypes of SAR11 [26]. An alternative hypothesis is that abundant variants of HTVC010P exist which are ecotype-specific, but which have not yet been captured in pelagiphage cultures, causing reads to recruit to the most closely related strain in the databases [23]. Viral single-amplified genomes (vSAGs) and metagenomically assembled viral genomes (MAVGs) of predicted pelagiphages (without cultured representatives) suggest that current viral isolates only cover a fraction of the available pelagiphage diversity [30, 35, 36].

In this study, we isolated and sequenced eleven strains of a novel pelagiphage species, *Pelagibacter* phage Skadi, named after the Norse goddess of winter. We demonstrate that Skadi is closely related to the globally abundant HTVC010P, both in gene sharing networks and single-gene phylogeny. Together with our previously isolated *Pelagibacter* phages Greip and Lederberg, as well as seven other HTVC010P-type pelagiphages, they form a distinct novel viral family, for which we propose the name *Ubiqueviridae*. Recruitment of reads from global marine viromes revealed that Skadi is a polar pelagiphage ecotype that dominates Arctic and Antarctic marine viral communities, but read recruitment rapidly decreases in viromes from lower latitudes. At tropical and subtropical latitudes, Skadi is near or below limits of detection, apart from regions with direct influx of polar water, such as the Humboldt Current System in the Southern Pacific Ocean. The global distribution pattern, in addition to experimentally confirmed host ranges, suggest that abundant populations of Skadi replicate in polar waters on cold-water ecotypes of SAR11 and are potentially transported around the globe in currents of sinking polar water which are part of the ocean conveyor belt. Skadi therefore serves as an exemplar of niche specialisation in pelagiphages and a model organism for determining how ocean currents might shape regional viral communities through the viral seed bank.

## Results & Discussion

### Skadi is closely related to Pelagibacter phage HTVC010P

A total of eleven Skadi-like (Average Nucleotide Identity (ANI) > 99 %) *Pelagibacter* phages were isolated on the SAR11 cold-water ecotype *Pelagibacter ubique* HTCC1062 from three coastal surface water samples taken from the Western English Channel (WEC) in September, October and November of 2018 [26]. Ten out of eleven genomes were assembled into single circular contigs, with one assembly returning a fragmented, linear genome. Four out of the ten circular contigs contained a direct terminal repeat of 125bp at the end of the linearized genome indicative of *pac* headful packaging similar to the T7-like P-SSP7 cyanophage [37]. General features of the genomes are shown in Table 1. Genome sizes ranged from 34,897 base pairs (bp) to 35,392 bp with a G+C content between 31.49 % and 32.19 %. All full-length genomes contained 65 putative open reading frames (ORFs) (Figure 1). Pairwise average nucleotide identity (ANI) values between strains ranged from 99.27% to 99.81%, all strains were therefore considered to be the same viral species [38], hereafter referred to as Skadi, unless a specific strain is specified. Skadi was related to Pelagiphage HTVC010P (ANI 32.9%) and other HTVC010P-type pelagiphages, based on whole-genome phylogenetic analysis for all *Pelagibacter* phages (Figure 2), but had a smaller capsid (39±0.7 nm (Figure 1B), compared to 50±3 nm for HTVC010P [22]). The difference in capsid size reported for HTVC010P [22, 23] and here for Skadi may be partly due to different operators and protocols for performing transmission electron microscopy (TEM). Like HTVC010P, the short stubby tail of Skadi suggested a *Podoviridae* morphology.

**Table 1.** Overview of general features and accession numbers for Skadi-like viral isolate genomes.

| Phage | Collection number | Genome Size (bp) | G+C (%) | Terminal Repeat (bp) | ANI (%) to Skadi-1 | ORFs | Accession number | Collection Date |
| --- | --- | --- | --- | --- | --- | --- | --- | --- |
| Skadi-1 | EXVC101P | 35,392 | 31.53 | 125 | 100 | 65 | OP131302 | Sept 24 2018 |
| Skadi-2 | EXVC102P | 35,392 | 31.49 | 125 | 99.763 | 65 | OP131299 | Sept 24 2018 |
| Skadi-3 | EXVC103P | 35,083 | 31.5 | n.a. | 99.506 | 65 | OP131297 | Sept 24 2018 |
| Skadi-4 | EXVC104P | 34,897 | 31.51 | n.a. | 99.274 | 65 | OP131301 | Sept 24 2018 |
| Skadi-5 | EXVC105P | 35,374 | 31.54 | 125 | 99.633 | 65 | OP131293 | Sept 24 2018 |
| Skadi-6 | EXVC106P | 35,392 | 31.57 | 125 | 99.811 | 65 | OP131294 | Sept 24 2018 |
| Skadi-7 | EXVC107P | 35,247 | 31.56 | n.a. | 99.638 | 65 | OP131300 | Nov 05 2018 |
| Skadi-8 | EXVC108P | 35,276 | 31.53 | n.a. | 99.554 | 65 | OP131296 | Nov 05 2018 |
| Skadi-9 | EXVC109P | 35,276 | 31.53 | n.a. | 99.554 | 65 | OP131303 | Nov 05 2018 |
| Skadi-10 | EXVC110P | 35,267 | 31.5 | n.a. | 99.813 | 65 | OP131295 | Nov 05 2018 |
| Skadi-11 | EXVC111P | 9,865 | 32.19 | n.a. | n.a. | n.a. | OP131298 | Sept 24 2018 |

**Figure 1.**
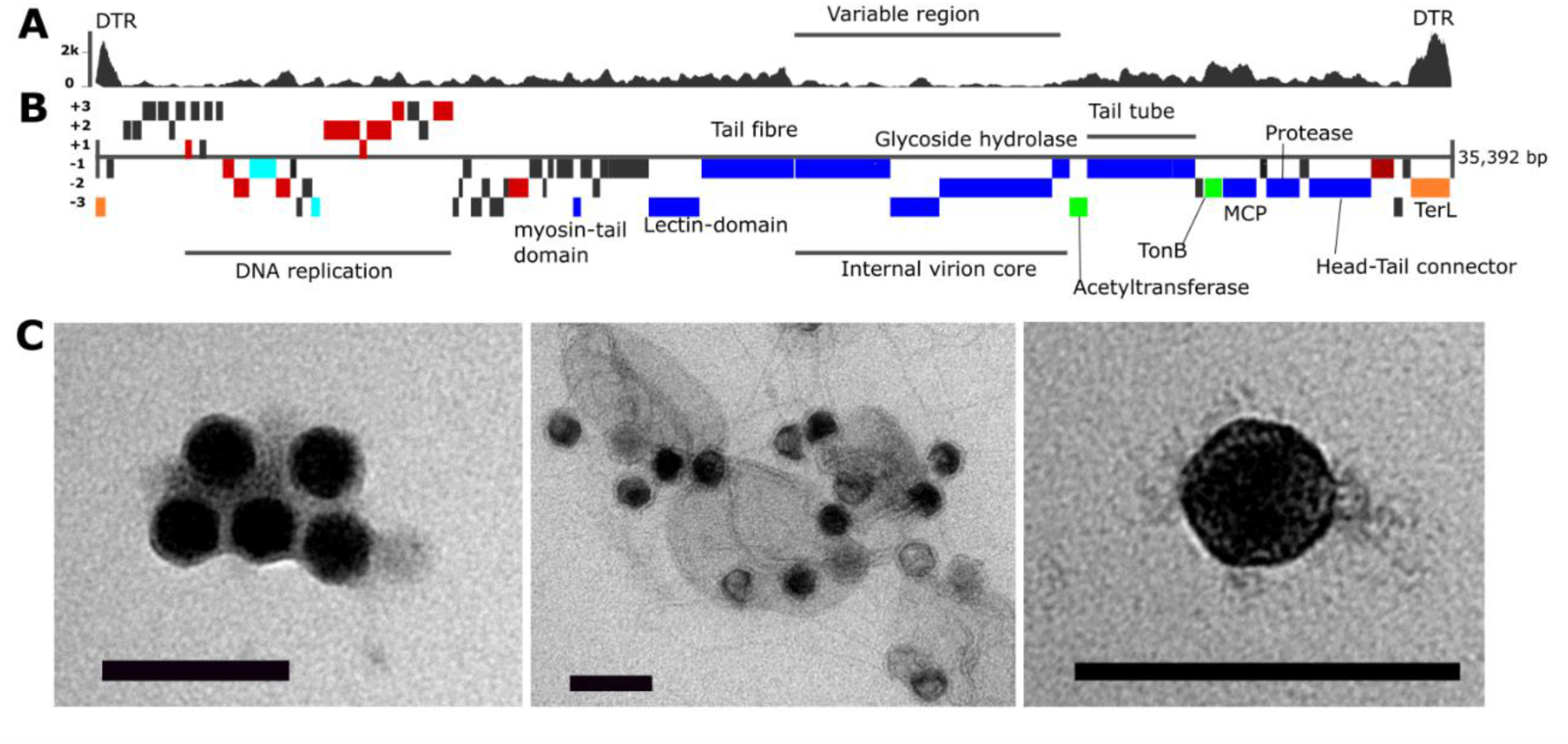
General features of *Pelagibacter* phage Skadi. **(A)** Plot showing per nucleotide coverage of the Skadi genome, high coverage at contig-ends suggest direct terminal repeats; the region encoding for virion proteins has low average coverage, suggestive of a hypervariable region **(B)** Genome map of Skadi, translation frames of Open Reading Frames (ORFs) are indicated in the figure, functional annotations of ORFs are color-coded with red: DNA replication and metabolism; teal: transcription; blue: Virus structural genes; green: virulence related genes; Orange: packaging. **(C)** Transmission Electron Microscopy (TEM) images, black scale bar represent 100 nm, left: icosahedral viral capsids of Skadi with a size of 39± 0.7 nm; middle: Skadi viral particles and cellular debris of lysed HTCC1062 host cells; right: Skadi single viral particle.

**Figure 2.**
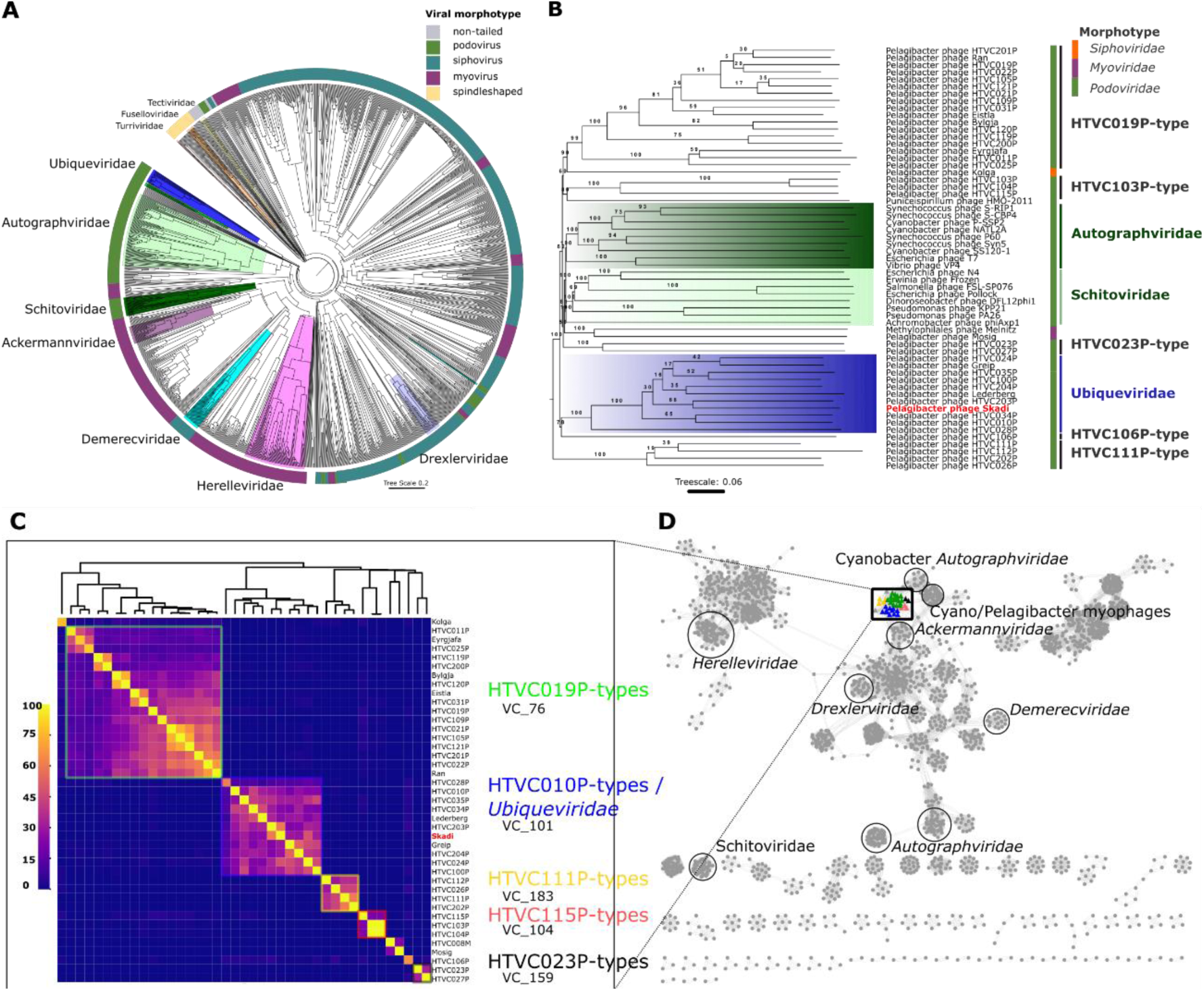
*Pelagibacter* phages can be grouped into distinct clusters. **(A)** All full length genomes were screened against DB-B:Baltimore Group Ib prokaryotic and archaeal, calculated using GRAViTy 1.1.0 (http://gravity.cvr.gla.ac.uk/). Outer ring represents the main virus morphotypes, branch highlights indicate ICTV recognised viral families. Blue highlighted branch represents the proposed *Ubiqueviridae* family. **(B)** Full-genome phylogenetic tree created in VICTOR [83] using known *Pelagibacter* phage genomes, leaf-label for Skadi is highlighted in red. Randomly chosen representative genomes of the *Autographviridae* (light green) and *Schitoviridae* (dark green) families were included and highlighted, the branch with all members of the proposed *Ubiqueviridae* is highlighted in blue. All bootstrap values are shown. **(C)** Heatmap of hypergeometric probability of shared genes between phages isolated on SAR11 hosts based on protein clusters identified by vConTACT2, highlights indicate corresponding vConTACT2 viral cluster assignments, no highlight indicates singleton viral clusters in vConTACT2 gene sharing analysis. Hypergeometric probability of shared genes between Skadi-like and HTVC010P-like phages suggests both are relatively closely related. **(D)** Visualised vConTACT2 analysis of shared gene network. The location of genomes of ICTV recognised bacteriophage families and genomes of phages infecting SAR11 with podovirus morphology are indicated, coloured nodes represent viral cluster identified by vConTACT2 and correspond to viral clusters colour coded in (A).

### Skadi uses host RNA and DNA polymerases for viral replication

Whole-genome alignment showed that the Skadi genome is arranged into early and late genomic regions similar to other HTVC010P-type genomes [27]. A TonB-dependent receptor protein located between the two tail tube proteins A and B and the major capsid protein suggests TonB as a putative receptor for adsorption and initiating infection [39, 40] (Figure 1). TonB-dependent transporters are one of the dominant proteins found in surface ocean metaproteomes, with increasing abundance in nutrient-rich, coastal waters [41]. We did not find Skadi-like TonB-receptors in other cultivated pelagiphages, suggesting that these species use different receptor proteins. We identified three internal virion proteins similar to gp14, gp15 and gp16 found in T7, which likely share a similar role in injecting phage DNA through the host periplasm during infection [42, 43]. A proximate glycoside hydrolase gene found in Skadi is likely a structural component of the internal virion complex, facilitating the injection of phage DNA by breaking glycosidic bonds in the cell wall, as was found to be the role of the glycoside hydrolase (Gp255) in *Bacillus* phage vB_BpuM_BpSp [44].

Skadi encodes for an acetyltransferase gene downstream of the internal core complex that may play a role in metabolic hijacking and host programming via host quorum sensing networks [45]. In a phiKMV-like virus infecting *Pseudomonas aeruginosa* acetyltransferases were responsible for the shutdown of host transcription by cleaving the bacterial RNA polymerase (RNAP) during the early stages of infection [46]. However, unlike pelagiphages in the *Autographviridae* family, Skadi does not encode a viral RNA polymerase (RNAP), making cleavage of host RNAP unlikely. Temporal regulation in the absence of endogenous RNAP is common in T4- and λ-like phages and involves sophisticated mechanisms that rely on early to late promoter sequences for recognition by the host RNAP [47]. We identified four s^70^ promoters and two Pribnow boxes (TATAAT-promoter sequences) in proximity of DNA replication and manipulation genes, but were unable to identify any terminator sequences, tRNA, tmRNA or ribonucleotide switches. This could suggest that Skadi uses host tRNAses, reinforcing similar codon usage between the host and phage, and potentially constraining host range [48]. Skadi also lacks a phage encoded DNA polymerase (DNAP), but encodes for two transcriptional regulator proteins, similar to HTVC010P-type phages [27]. Likely these transcriptional regulators are used to hijack the cellular transcriptional machinery, indicating that Skadi, as predicted in other HTVC010P-type phages, relies on host RNAP and DNAP proteins for translation and transcription.

An additional methylase gene likely provides Skadi with protection from host restriction endonucleases found in *Pelagibacter ubique* HTCC1062 (SAR11_RS00560), or might help to protect the Skadi virocells against superinfecting DNA from phage competitors as endonucleases are a common feature in pelagiphages [49]. Skadi further encodes peptide hydrolase and nuclease genes that are likely involved in breaking down host proteins and nucleic acids during the early infection stage to recycle material for phage synthesis [50]. Skadi encodes a helicase loader protein similar to the λ-like DNA replication machinery, without virally encoded genes for helicases or primases. In addition to the lack of a terminator sequence, this suggests that Skadi utilises ‘rolling-circle’-like DNA replication. Skadi encodes several common phage structure protein, namely: three internal virion proteins, major capsid protein, tail tube proteins A and B, tail fibre, head-tail connector and prohead-protease. Located within the direct terminal repeat region is a conserved TerL gene for packaging. For cell lysis, Skadi encodes a peptidase M15, similar to other HTVC010P-type pelagiphages [27].

### Skadi contains an arms-race associated hypervariable region that is conserved across HTVC010P-like phages

Alignment of HTVC010P-type genomes showed that regions related to DNA replication and metabolism, as well as the terminase and phage head related proteins are conserved, using recommended amino acid identity (AAI) thresholds of >30% [38] (Supplementary Figure 1). In contrast, the region of the genome encoding virion core proteins and tail fibre related genes had an AAI identity of <30% between HTVC010P-types and our phages. Previous analyses of long-read sequencing based viromes from the Western English Channel resolved the microdiversity of HTVC010P across niche-defining genomic islands, and revealed a hypervariable region of ∼5000 bp in length that contained a ribonuclease and an internal virion protein [51]. Such regions of low coverage occur when reads from a diverse population fail to recruit to the genome of a specific strain and are indicative of genetic population variance [52]. Mapping randomly subsampled (5 million reads per virome) from all environmental GOV2 viromes against HTVC010P-type genomes revealed an average per nucleotide coverage of 4,745 × for Skadi per sample. The region between 13,646 and 15,713 bp returned per nucleotide coverage of less than 100, marking it as a putative hypervariable region (HVR) using previously defined cut-offs (<20% of the median contig coverage and at least 500 kb in length) [53]. Five out of 11 other HTVC010P-like phages recruited at least the minimum number of mapped reads and coverage to putatively identify HVRs, and these genomes showed consistently low recruitment in the same genomic region as Skadi, showing this could be conserved feature within this group. Skadi’s putative HVR spanned two ORFs encoding an uncharacterized phage protein and a structural lectin-domain containing protein. Lectin-folds in association with phage tail fibre proteins are often involved in host receptor binding and cell attachment [54]. These proteins might also be used by Skadi to attach to SAR11 cells, supporting the idea that the hypervariable region is a hotspot for arms-race co-evolution and selection within global Skadi populations.

### *Skadi and other HTVC010P-type pelagiphages constitute a novel viral family within the* Caudovirales order

Analysis of average nucleotide identity between Skadi and all other known *Pelagibacter* phages showed that Skadi’s closest relatives were the phages Greip (ANI 37%) [26], Lederberg (ANI 33%) [25] and the highly abundant HTVC010P-type pelagiphages (ANI 30% to 49%) [22, 27]. The average nucleotide identity between all these phages was <60%, which, based on established ICTV thresholds, suggest that each phage are sole representatives of separate genera [38]. Other known pelagiphages with podovirus morphology had ANI values <5%. A phylogenetic tree based on complete genome-based proteomics placed Skadi and HTVC010P-like phages on a monophyletic branch distinct from other phage groups, providing support for a family-level taxonomic assignment (Figure 2). This comparative phylogenomic analysis against all dsDNA prokaryotic and archaeal viruses within the DB-B:Baltimore Group also suggested that there are no other cultivated phages in the database which are closely related to Skadi and HTVC010P-type phages. Full-length genome based phylogenetic analysis using 17 arbitrarily chosen representatives of the *Schitoviridae* and *Autographviridae* families, which are the two main ICTV recognised families with podovirus morphology, together with all 40 *Pelagibacter* phages also placed Skadi and HTVC010P-type phages on a well-supported branch distinct to all other branches (Figure 2B). This suggests that these phages are members of the same family-level taxonomic group, for which we suggest the name *Ubiqueviridae* – derived from the latin word *ubique* (“everywhere”) to reflect both the high abundance and ubiquitous distribution of these phages as well as in recognition of their common host, *Pelagibacter ubique*.

Further evidence that *Ubiqueviridae* represents a distinct taxonomic cluster from other pelagiphages was provided by hypergeometric probability of shared protein clusters (Figure 2C and D), showing high probability of shared genes within groups corresponding to pelagiphage types, with *Ubiqueviridae* forming a single viral cluster. The taxonomic assignment tool vConTACT2 placed the *Ubiqueviridae* into a separate viral cluster from other pelagiphages. Visualising a shared gene network analysis with all RefSeq phages showed that podophages infecting SAR11, as well as the pelagisiphophage Kolga, share genes between pelagiphage viral clusters at a higher level compared to pelagiphages and viruses of other hosts. PIRATE analysis [55] identified ten shared core genes within all eleven members of the proposed *Ubiqueviridae* family (∼16% of ORFs in Skadi) (Supplementary Table 1). *Ubiqueviridae* did not share core genes with pelagiphages of the HTVC019P-type genomes (Unconfirmed part of the *Autographviridae* family) or taxonomically unclassified HTVC111P-types, HTVC103P-types or HTVC023P-types. One shared core gene (*TerL*) was found between *Ubiqueviridae* and the singleton HTVC106P. Phylogenetic analysis of *TerL* shows that the *Ubiqueviridae* genes form a monophyletic branch, with HTVC106P as a closely related but separate branch (Supplementary Figure 2). Additional phylogenetic analysis on individual structural genes (tail tube and major capsid proteins) that are commonly found in dsDNA podophages also supports the proposed *Ubiqueviridae* as a monophyletic group and places other pelagiphage types onto distinct and well-supported (bootstrap value > 0.94) monophyletic branches (Supplementary Figures 3 and 4). Other pelagiphage types had 15 to 33 shared core genes in their respective groups (Supplementary Table 2). No core genes were shared between *Ubiqueviridae* and siphophages (*Pelagibacter* phage Kolga), nor with myophages (*Pelagibacter* phages Melnitz and HTVC008M) that have been experimentally confirmed to infect SAR11 [22–24, 27, 56].

### Global distribution patterns reveal Skadi as a polar surface pelagiphage ecotype

Skadi was observed almost exclusively in high latitude surface waters in both the Arctic and Southern Ocean, despite the fact that both bodies of water are separated by the Atlantic and Pacific Ocean, respectively (Figure Skadi 3A). Skadi was near or below limits of detection in viromes from lower latitudes (below 66° absolute latitude). Based on the absence of Skadi populations at lower latitudes, we speculate that it is unable to replicate sufficiently to maintain sizable populations in these waters. Therefore, the Atlantic and Pacific Oceans form an evolutionary barrier separating southern and northern populations. As ^14^C carbon isotope analysis has shown that a single “parcel” of seawater can take up to 1,000 years to be transported around the world by global ocean circulations [57], the high abundance of genetically similar Skadi populations in both polar regions (based on read recruitment) suggests that they form conserved populations despite the implied relatively low rates of direct genetic exchange.

To further evaluate differences in Skadi populations, we applied Single Nucleotide Variant (SNV) profiles [21] using 22 GOV2 metagenomes in which Skadi was sufficiently abundant to meet a minimum coverage cutoff of 70 %. In total, 10,142 (461 SNV positions across 22 metagenomes) entries were evaluated (91,410 were removed, following recommended cutoffs for coverage and minimum departure constraints [58]). These 461 positions spanned 13 of the 60 Skadi genes. Population structures were evaluated using clustering of SNVs identified in each metagenome and revealed two groups (Supplementary Figure 5): The first group comprised 12 samples obtained from latitudes greater than 60° N (considered here as polar influenced biomes), as well as two samples (sample 82, surface and DCM) obtained from 47° Southern Atlantic off the Argentinian coast ([FKLD] Southwest Atlantic Shelves Province). The second group comprised 10 samples retrieved from tropical and subtropical samples from the South Atlantic, South Pacific and the Mediterranean, spanning between latitudes 34.95° S and 42.17° N. Metagenomes from different depths but same locations clustered tightly together except one (25_SRF and 25_DCM), suggesting depth and associated intrinsic changes in physico-chemical conditions is not a factor that differentiate Skadi populations, compared to geographical location. Skadi was virtually absent from metagenomes associated with the North Pacific Subtropical Gyre, with only three out of 382 metagenomes (SRR9178452, SRR9178352, SRR9178296 [59]; all from 200m and 250m depth) from Station ALOHA (22°45′ N, 158° W) meeting minimum coverage cutoff requirements. This suggests that cold-(deep-)water at lower latitudes does not support large Skadi populations.

Reported mutation rates in dsDNA viruses range from 10^−4^ to 10^−6^ substitutions per site per cell infection [60–62]. Assuming Skadi latent period is comparable to the latent periods of 22-24 hours reported for HTVC010P [22], and considering Skadis 35392 bp genome, a rough estimate of 0.04 to 3.54 substitutions per year would be expected over the full genome, or 35.39 to 3539 substitutions in the 1,000 years that it can take to transport water around the world by ocean circulations [57]. Therefore, we propose that the barrier formed by the Atlantic and Pacific Oceans does not create genetically distinct Skadi populations at either polar region, suggesting that ocean mixing occurs faster than Skadi mutation rates.

Based on the study of T4-like marine cyanophages, it has been suggested that the abundance of virus populations reflects that of their hosts [15]. Pelagiphage HTVC010P was initially isolated from the subtropical Sargasso Sea on cold-water ecotype *Pelagibacter ubique* HTCC1062 [22], which decreases in a north-south gradient to be near-absent at low latitudes, where HTCC7211 is highly abundant [63, 64]. We used competitive recruitment of short reads from marine viromes (GOV2) to evaluate relative abundances of all 41 isolate genomes of pelagiphages across global oceans (≥95% nucleotide identity over 90% of read length, with 40% minimum genome coverage threshold to reduce false positives [32]). HTVC010P did not have any discernible spatial distribution patterns and recruited a broadly uniform number of reads across all latitudes (Figure 3B). Therefore, we speculate that HTVC010P can infect both warm- and cold-water ecotypes, possibly with lower efficiencies in cold-water ecotypes, explaining its abundance and uniform distribution across the global oceans, and its replacement by Skadi at higher latitudes. As with pelagiphage HTVC010P, Skadi and Greip were isolated using the SAR11 cold-water ecotype HTCC1062, but when these phages were used to challenge two warm-water ecotypes which failed to produce a reduction in host numbers, indicating preferential infection of the cold-water host ecotype. Low abundance of Skadi at temperate and tropical latitudes is most likely explained by a lack of suitable hosts for efficient replication (Figure 3C), highlighting how phage ecotypes are constrained by environmental selection of their preferred hosts, similar to observations made in cyanobacteria and roseobacteria virus-host systems [65, 66].

**Figure 3.**
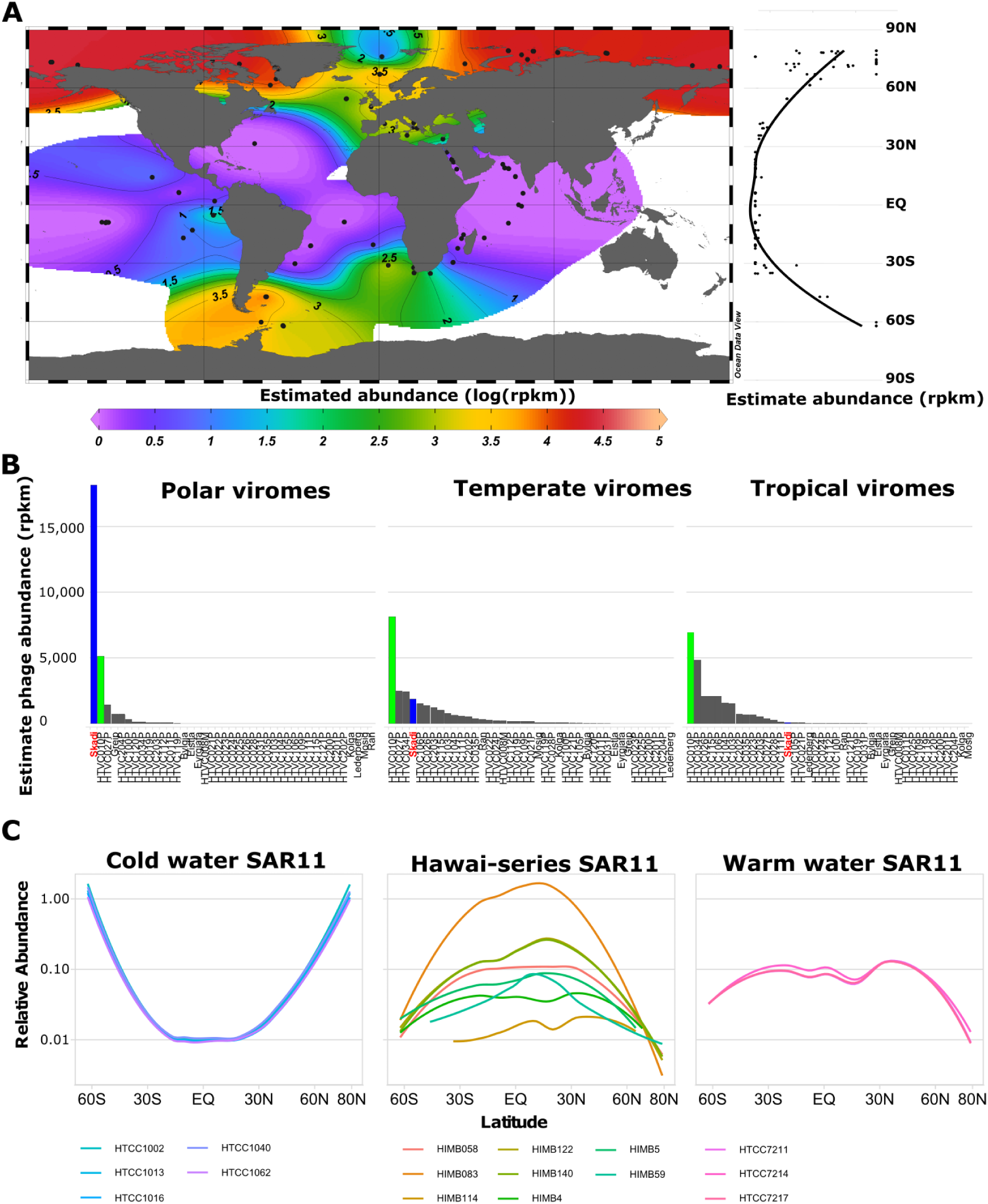
Global distribution pattern of *Pelagibacter* phage Skadi. **A** Estimated abundance in reads per genome kilobase mapped per million reads (RPKM) of Skadi’s full length genome from the GOV2 dataset; **B** Estimated abundance (RPKM) of isolated phages infecting SAR11 (blue: Skadi; green HTVC010P) grouped by absolute latitude into “Polar” (90° to 66°), “Temperate” (66° to 33°) and “Tropical” (33° to 0°) regions; **C** Relative abundance of SAR11 bacterial strains based on single amino acid variants from Delmont et al. [21].

Across all GOV2 samples, global median relative abundance of HTVC010P (3100 - 7713 RPKM) was significantly higher than that of Skadi (0 – 1673 RPKM) by an average effect size of 5,141 RPKM (2699-7530 RPKM). Repeating this analysis with mean RPKM (sensitive to outliers) instead of median RPKM removed this significant difference, as expected given the highly skewed abundance of Skadi within polar regions. The mean difference between Skadi and HTVC034P (the next highest recruiting pelagiphage) was 3,902 RPKM (2088 – 5776 RPKM), indicating that Skadi and HTVC010P are the two most abundant pelagiphages in the oceans by ∼2-20-fold. Mean relative abundance of the 39 remaining pelagiphage ranged from 0 - 2,562 RPKM, suggesting that global oceans are dominated by few highly successful pelagiphages and a long tail of less abundant variants. This supports the viral Seed-Bank hypothesis for SAR11 virus-host systems, which suggests that marine viral genotypes are ubiquitously distributed albeit relatively rare, but can become abundant when environmental conditions enable efficient replication in available suitable hosts [33].

Relative abundance of Skadi was significantly negatively correlated to temperature (RPKM∼temperature linear regression, p < 0.001, R^2^=0.69) (Supplementary Figure 6). In contrast, relative abundance of HTVC010P and the majority of other pelagiphages (21 out of 28 pelagiphages with non-zero RPKM in >3 viromes) was not significantly correlated to temperature (p= 0.06, R^2^=0.08 for HTVC010P, R^2^ <0.5, p≥0.05 for other pelagiphages). *Pelagibacter* phage HTVC027P was the only phage other than Skadi to have a significant, but weak, negative correlation with temperature (p < 0.001, R^2^=0.27). *Pelagibacter* phage HTVC023P significantly and positively correlated with high temperatures (p=0.003, R^2^=0.59), suggesting an association with warm-water SAR11 ecotypes, though the original isolating host was the cold-water HTCC1062, indicating potential generalism in HTVC023P, similar to that predicted for HTVC010P. This is in line with two long-term 16S rRNA time series conducted at BATS station in the Sargasso Sea and L4 station in the WEC, which found that SAR11 Ia3 have a broad temperature at which it can thrive (from 7 to 30 °C), whilst the maximum temperature for Ia.1 was ∼20 °C [67]. This suggests that phages specialising for SAR11 Ia.3 hosts are less likely to display significant trends with temperature, whereas abundance of phages specialising on cold-water SAR11 Ia.1 would be expected to inversely correlate with temperature. A similar temperature dependent relationship between viruses and environmental parameters has recently been demonstrated for HMO-2011-type phages infecting Roseobacteria [66]. Our results indicate a similar ecotypical distribution is present for pelagiphages, suggesting that this is a common occurrence for viruses of marine heterotrophs.

Given the importance in sinking polar water driving large-scale global water circulation [68, 69], we evaluated whether Skadi may also be found in lower latitude viromes known to be under the influence of deep-water upwelling. An intriguing exception to the otherwise polar distribution pattern for Skadi was found in three GOV2 samples (100_DCM, 102_DCM, 102_MES), from which Skadi recruited between 517 and 912 RPKM. These samples were taken in the eastern parts of the Pacific Equatorial Divergence Province (PEOD) and the South Pacific Subtropical Gyre Province (SPSG) in autumn 2011 [70]. An important feature of this region is the Humboldt Current System (HCS), which is a cold-water current flowing from Antarctica in parallel to the South American west coast. This causes upwelling of nutrients along the coastline, creating one of the most productive regions in the ocean, but also feeds cold water from the Southern Ocean into the Pacific Ocean [71, 72]. We hypothesize that dominant Skadi populations in the Southern Ocean were transported via the HCS into the Southern Pacific Ocean, where over time Skadi populations are replaced by other actively replicating phages like HTVC010P as the most dominant pelagiphages (4,829 to 5,191 RPKM). As Skadi was only the 8th to 14th most abundant pelagiphages in the Southern Pacific (compared to the Southern Ocean where Skadi was the most abundant pelagiphage) it is likely that Skadi populations are either outcompeted by phages that are better adapted to local conditions, or that regional SAR11 populations do not provide enough suitable host cells for Skadi to maintain a dominant population. In this scenario, if the population dynamics conform to the Royal-Family hypothesis [16], the Southern Pacific Skadi population is likely in a state of decay and being succeeded by the closely related HTVC010P.

Abundance of *Pelagibacter* phage Greip, previously identified as a potential polar ecotypic phage [26], did not significantly correlate to temperature (p=0.38, R^2^=0.08), but was detected in five viromes taken in the Arctic with temperatures of <5 °C and only two viromes from below 66° north and south. Greip was isolated through enrichment on HTCC1062 in the WEC when ambient surface temperature was 14.8 °C. However, Greip was below the limit of detection in a WEC virome [51]. Similarly, Skadi was isolated on eleven separate occasions from the WEC, with water temperatures ranging from 14.1 to 15.5 °C, but was present in a WEC virome (5300 RPKM). This could suggest that unlike Greip, Skadi is able to replicate on local SAR11 populations in this environment, which indicates different optimal hosts for Greip and Skadi, despite both of them being able to replicate on HTCC1062. Successful propagation of Greip and Skadi in cultures kept at 15 °C further indicates that host-availability, rather than temperature itself, limits viral replication in these putative viral polar ecotypes.

### Ecotypic pelagiphages are an important aspect of global viral communities

Cold-water environments are often defined as areas with average annual temperatures below 15 °C [73], with polar oceans temperatures between +5 °C and -2 °C, which allows for up to double the available oxygen concentrations compared to water at 20 °C [74]. The Arctic Ocean also experiences high riverine influx of terrestrial nutrients [75], and subsequently high primary production [76] (for which chlorophyll *a* serves as a proxy). Constrained ordinations provide further evidence that Skadi abundance is correlated with high latitude and/or associated low temperature (Figure 4), but also suggest that high oxygen and increased concentration of nutrients and chlorophyll *a* correlate positively with Skadi abundance. Absolute longitude did not impact ordinations for any pelagiphage, indicating that distribution patterns are a function of shared environmental conditions associated with geographical provinces, but not the marine geographical provinces themselves. This is in line with our observations of Skadi populations at each pole that are putatively connected via the global conveyor belt, as ocean currents would continuously mix global viral communities, maintaining the viral Seed-Bank. These results therefore support Skadi as a polar specialist. High temperatures correlated positively with relative abundance of five other (lower abundance) pelagiphages in addition to HTVC023P (HTVC103P, HTVC104P, HTVC115P, HTVC106P, HTVC026P). However, 18 out of 41 pelagiphage genomes did not recruit reads from enough viral metagenome samples (non-zero RPKM in five or more metagenomes) to establish any robust patterns, indicating that these phages are part of the respective Seed-Bank at sampling locations. Considering that the Seed-Bank hypothesis addresses the turnover of dominant phages, the fact that across space and time both HTVC010P and Skadi remain dominant, suggests that the aforementioned ecological constraints maintain the host population and subsequently the dominance of a small group of phages – supporting the Royal-Family hypothesis [16]. Deep-sequencing of viral *phoH* genes in the Sargasso Sea also suggested that the majority of operational taxonomic units (OTUs) remained rare throughout the sampling period, with a small number of OTUs dominating throughout the seasons, depths and years [77]. Similarly, in SAR11 virus-host systems our results could mean that abiotic factors shaping SAR11 ecotypic niche partitioning also indirectly maintain the dominance of a small number of phages out of a large diverse pelagiphage community akin to pelagiphage ecotypes.

**Figure 4.**
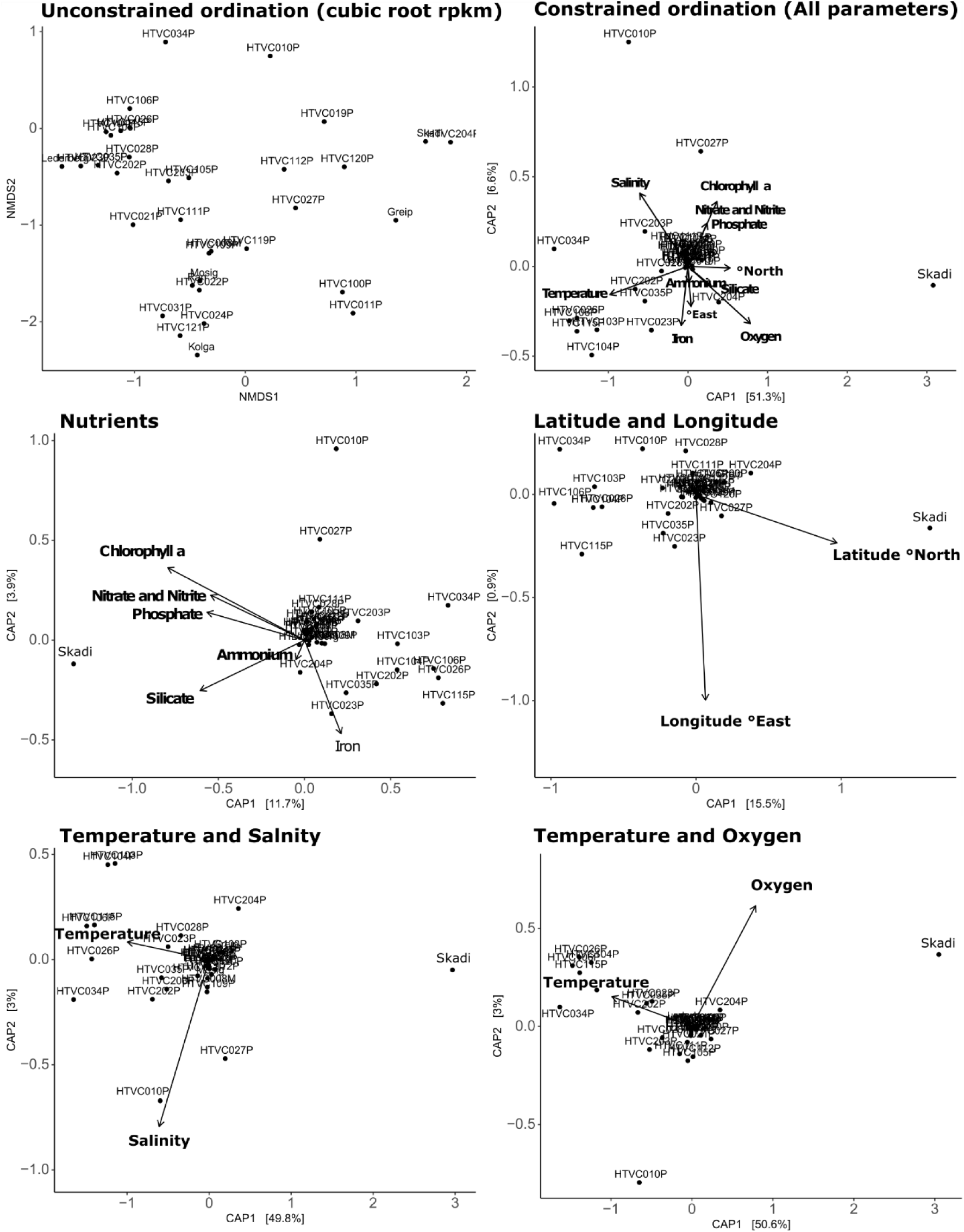
Skadi’s abundance is driven by typical cold-water parameters. Plot shows ordinations of RPKM values obtained from GOV2 viromes constrained with different environmental parameters provided by the GOV2 metadata. Skadi abundance is driven by low temperature, high latitude and high nutrient concentrations, HTVC010P abundance is driven by low oxygen and high salinity, but not temperature

## Conclusion

In this study we isolated and characterized the highly abundant pelagiphage Skadi, defining it as one of the most abundant phages infecting the ubiquitous SAR11 clade. Phylogenetic analysis and comparison to other pelagiphages shows that Skadi and HTVC010P-type phages are a monophyletic and distinct taxonomic unit for which we propose the creation of a novel viral family named *Ubiquiviridae*. Using Skadi and HTVC010P as examples, we show that both putative generalist and specialist pelagiphages can be successful and dominant. Our metagenomic analysis of the GOV2 viromes shows that pelagiphage distribution is ubiquitous as thought previously, but the majority of pelagiphage species remain in the low abundance fraction, conforming to the Seed-Bank model. Skadi abundance is strongly correlated to polar environmental conditions, marking it as a pelagiphage cold-water specialist. The high abundance of Skadi at both poles further suggests that SAR11 ecotypic niche partitioning indirectly shape the pelagiphage community akin to pelagiphage ecotypes, but also that the dominance of these viral species is being maintained across space and time as suggested by the Royal-Family model. Furthermore, considering the virome sampling locations across global oceans, observations of increased abundance for Skadi populations along the Humboldt Current Systems demonstrates how ocean currents could potentially maintain the viral Seed-Bank globally. If polar oceans keep increasing in temperature due to escalating climate concerns, a shift in the polar microbial community is likely [78]. By uncovering polar ecotypic distribution of an abundant pelagiphage, our results suggest that polar pelagiphage communities could be altered under predicted warming patterns, with polar specialists lost due to the inability of further poleward niche expansion. SAR11 ecotypes are known to possess specific preferences for different carbon moieties, with different pathways that produce climactically active gasses [19]. Coupled with the importance of viral predation on community productivity and carbon export from the surface ocean [79], it is possible that this loss of viral diversity and shifting of virus-host dynamics may influence biogeochemical cycles.

## Methods summary

Methods deployed in this study are described in detail in the supplementary materials. Briefly, all phages were isolated on host *Pelagibacter ubique* HTCC1062 using Dilution-to-Extinction methods with surface water samples from the Western English Channel (50°15.00 N; 4°13.00 W) as described previously [26]. Viral particles were precipitated using a modified PEG8000/NaCl DNA isolation method [80] and purified with Wizard DNA Clean-up kits (Promega), following the manufacturer’s instructions. Sequencing of phage DNA was performed using Illumina 2 × 150 PE sequencing. Genomes were annotated and manually curated using an approach developed for the SEA-PHAGES program [81], where the output from multiple annotation programs is imported into DNA Master (v5.23.3) and evaluated by using a scoring system to minimize human bias. Phylogenetic analysis was performed using ICTV recommended tools and cut-offs for viral phylogeny [38]. To estimate global relevance of Skadi, sequencing reads were mapped against the GOV2 database [82] and viromes from the ALOHA station time series (22°45′ N, 158° W) in a subtropical Pacific ocean gyre [59].

## Data availability

Annotated genomes of the phage isolated in this study are deposited in NCBI’s GenBank under accession number OP131293-OP131303, sequencing data can be found in BioSample submissions SAMN30128164-SAMN30128174. All submissions are associated with BioProject PRJNA625644.

## Acknowledgements

We thank Christian Hacker and the Bioimaging Centre of the University of Exeter for performing TE microscopy and imaging. We would also like to thank the crew of the R/V *Plymouth Quest* and our collaborators at the Plymouth Marine Laboratory for providing water samples. We acknowledge the IT team at the University of Exeter and using the High-Performance Computing (HPC) facilities. Phage genome sequencing was provided by the Exeter Sequencing Service. Holger H. Buchholz was funded by the Natural Environment Research Council (NERC) GW4+ Doctoral Training program. LMB and BT were funded by NERC (NE/ R010935/1) and the Simons Foundation BIOS-SCOPE program.

