## Supplementary Materials for "*Pelagibacter* phage Skadi - An abundant polar specialist that exemplifies ecotypic niche specificity among the most abundant viruses on Earth"

*SAR11 hosts and growth conditions*

The SAR11 strains *Pelagibacter* HTCC1062 and HTCC7211 were kindly provided by the Giovannoni lab (Oregon State University, USA) and grown in the Artificial Seawater Medium (ASM1) [28] amended with 1 mM $NH_4Cl$, 10 µM $KH_2PO_4$, 1 µM $FeCl_3$, 100 µM pyruvate, 25 µM glycine, 25 µM methionine as well as 1 nM HMP, pantothenate, Biotin, Pyrroloquinoline quinone PQQ and Vitamin B12 each [29]. Continuous cultures were cultivated in 50 mL acid-washed (10% HCl) polycarbonate flasks at 15 °C without shaking.

*Water sampling and viral isolation*

1 L of seawater was collected in rosette-mounted Niskin bottles at a depth of 5 m from the Western English Channel Observatory coastal station L4 (WCO; http://www.westernchannelobservatory; 50°15.00N; 4°13.00W) on the 2018-09-24. Seawater was transferred immediately to a clean 2 L acid-washed polycarbonate (PC) Nalgene bottles (Thermo Fisher Scientific, Waltham, USA) and placed in a cooler box (Igloo, Katy, USA) at ambient temperature. Upon return to shore, water was transported to the University of Exeter for immediate processing (Two hours maximum duration from collection to processing). The sample was then processed and used for viral isolation on the host strain HTCC1062 as described previously [26]. Briefly, larger plankton was removed from the sample using a series of filter (GF/D 2.7 µm filters, 142mm 0.2 µm PC filter), and concentrated to ~50 mL with a 50R VivaFlow tangential flow filtration unit with a 100 kDa Hydrosart membrane (Sartorius instruments, Goettingen, Germany). The concentrate was filtered through a 0.1 µm pore PC syringe filter to remove remaining small bacteria, and used as inoculum (10% v/v) to infect exponentially growing HTCC1062 cultures in 96-well Teflon plates (Radleys, UK), prepared with ASM1. Growth of all wells was monitored using standard flow cytometry on 20 µL sub-samples diluted in 180 of µL 1x PBS, stained with SYBR Green. No-virus controls were compared to virus-treated samples to spot lysis. Cells in wells where lysis was observed were removed using 0.1 µm pore PC syringe filters, and used to prepare a dilution-to-extinction series in 10-fold dilution steps, that were then used to infect freshly prepared host cultures in 96-well Telfon plates, prepared as described above. This process was repeated three times, before a viral culture was considered ready for downstream applications.

*Transmission Electron Microscopy of viral isolates*

For ultrastructural analysis, virus particles were transferred onto pioloform-coated electron microscopy (EM) copper grids (Agar Scientific, Standsted, UK) by floating the grids on droplets of virus-containing suspension for 3 min. Following a series of four washes on droplets of deionized water, the bound virus particles were contrasted with 2 % (w/v) uranyl acetate in 2 % (w/v) methyl cellulose (mixed 1:9) on ice for 8 min and the grids then air-dried on a wire loop after carefully removing excess staining solution with a filter. Dried grids were inspected with a JEOL JEM 1400 transmission electron microscope operated at 120 kV and images taken with a digital camera (ES 1000W CCD, Gatan, Abingdon, UK).

*DNA isolation and sequencing*

HTCC1062 cultures in 300 mL ASM1 medium (prepared as described above) were grown in 1 L PC flasks at 15 °C without shaking. When cell density reached $1 \times 10^6$ cells/mL, 30 mL of viral lysate were added. Infected cultures were incubated for about one week. The cultures were transferred were transferred to 50 mL falcon tubes and centrifuged for 120 minutes (GSA rotor, Thermo Scientific 75007588) at 8,500 rpm/10,015 × g to remove larger cellular debris. To remove remaining cell fragments, supernatant was filtered through pore-size 0.1 µm PVVDF syringe filters. Viral particles were precipitated using a modified PEG8000/NaCl DNA isolation method [80]. Briefly, per 50 mL lysate, 5 g PEG8000 and 3.3 g NaCl were dissolved using Falcon tubes and incubated on ice overnight. The precipitate was pelleted by centrifugation at 8,500 rpm/10,015 × g for 90 min at 4 °C. After discarding the supernatant, the pelleted virus particles were re-suspended by rinsing the tubes twice with 1 mL SM buffer (100 mM NaCl, 8 mM $MgSO_4 \cdot 7H_2O$, 50 mM Tris-Cl). DNA was cleaned and extracted using the Wizard DNA Clean-Up system (Promega) following manufacturer's instructions, and eluted from the resin-columns using nuclease free water (60 °C). DNA libraries were prepared and sequenced using the services of MicrobesNG (Birmingham, UK), opting for standard Nextera protocols, targeting minimum 30-fold coverage with Illumina paired-end [2 × 250 bp] sequencing on the HiSeq 2500. Raw reads were trimmed, quality controlled and error corrected using bbmap and tadpole [84]. Viral genomes were assembled using SPAdes v3.13 [85] and evaluated with QUAST [86]. The trimmed reads were mapped back against the contigs for scaffolding using bowtie2 [87], BamM (alignment 0.9, identity 0.95) (available at https://github.com/minillinim/BamM) and samtools [88].

*Genome annotation*

Viral contigs were confirmed with VirSorter (categories 1 or 2, >15kbp) and only accepted if the contig showed mean depth coverage an order of magnitude greater than the next highest recruiting contig (to filter out any cellular DNA carryover). Gene calls returned by VirSorter were imported into DNA Master for manual assessment and curation [81]. Additional gene calls were made using GenMark [89], GenMarkS [90], GenMarkS2 [91], GenMark.hmm [92], GenMark.heuristic [93], Glimmer v.3.02 [94], and Prodigal v.2.6.3 [95]. All gene calls were tabulated and compared using a scoring system which evaluates gene length, gene overlap and coding potential of ORFs [81]. ORFs were then annotated using BLASTp against the NCBI's non-redundant protein sequences [96], Phmmer [97] against UniProts UniProtKB and uniprotrefprot [98], Swissprot [99] as well as InterProScan [100] and Pfam [101]. Virally-encoded tRNAs were searched for with the web applications of tRNAScan-SE v2.0 [102] and ARAGORN [103]. Genomes were scanned for riboswitches using the web application of RiboswitchScanner [104]. FindTerm (energy score < -11) and BPROM (LDF > 2.75) from the Fgenesb_annotator pipeline [105] were used to predict promoter and terminator sequences, using default parameters. Promoter sequence 5'-TATAAAT-3' [106, 107] and MotA box (TGCTTtA) dependent promoters were predicted by aligning these sequences against the whole genome using a BLASTn search. Predicted promoter/terminator sequences that were intergenic or within 10 bp of the start/stop of

ORFs were disregarded. Direct terminal repeats were identified with CheckV [108] (Default settings) and confirmed through a doubling of read coverage at contig ends [109].

*Phylogenetic analysis*

For the phylogenetic analysis we followed the roadmap for phage taxonomy [38], which was based on the most recent ICTV guidelines for viral taxonomy [110]. Species level genome comparisons were done using VIRIDIC v1.0 [111] with a species similarity threshold of 95% and genus threshold of 70%. Whole-genome based phylogenetic trees were constructed with VICTOR (formula $d_6$) through the online application available at https://ggdc.dsmz.de/victor.php [83]. Additionally, a proteomics tree for Pelagibacter phages against DB-B:Baltimore Group Ib prokaryotic and archaeal was created using GRAViTy 1.1.0 (accessed via http://gravity.cvr.gla.ac.uk/, 2 June 2021). For the shared genes of all 41 known pelagiphage isolate genomes were called using default prodigal v2.6.3 [95] and imported into the Discovery Environment 2.0 at https://de.cyverse.org/, where vContact-Gene2Genome 1.1.0 was used to prepare protein sequences for analysis with vConTACT2 (v.0.9.8.) in default settings [112]. Cytoscape v3.7.2 [113] was used to visualize the results from vConTACT2. Single-gene based phylogeny was done within the phylogeny.fr webserver [114], opting for default MUSCLE alignment [115] and the built-in curer removing only positions with gaps to conserve more positions for constructing phylogenetic trees. For the phylogenetic tree building, maximum likelihood trees were constructed with PhyML (100 bootstraps, unless stated otherwise) [116, 117] and visualized using FigTree (v1.4.4 available at http://tree.bio.ed.ac.uk/software/figtree/). Prokka v1.14.6 [118] in standard settings was used to create genome feature files (gff) of all Pelagibacter phages with podophage morphology, used as input for PIRATE v1.0.4 [55] for identifying orthologous core genes shared between and within family level groups using 'PIRATE --steps "30,40,50,60,70,80,90" -k "--e 1E-5 --hsp-length 0.5" -r –a' flags based on ICTV recommended thresholds of >30% identity and >50% coverage for constructing a pangenome [38]. To identify variable regions in the genome, bowtie-2 (v.2.3.5.1) [87] and sorted with samtools (v.1.11) [88] were used to map reads from the GOV2 dataset [82] with flags: 'bowtie2 –-seed 42 –-nondeterministic | samtools view –F 4 –bS' against full length genomes, which were rearranged so that the region encoding TerL was last in each genome respectively. Results were visualized with python 3.0 (www.python.org) and packages imported from the biopython project [119]. All figures were edited in Inkscape (www.inkscape.org) for aesthetics. Metagenomic reads were subsampled to 5 million reads for each virome using the reformat.sh within the bbmap suite. Bowtie2 v.2.3.5.1 [87] with the flags 'bowtie2 –-seed 42 –-nondeterministic' was used to created indexes from all known pelagiphage genomes [22–27] and sorted with samtools (v.1.11) using 'samtools view –F 4 –bS' flags [88].

*Metagenomic read mapping and global abundance analyses*

Global Oceans Viromes (GOV2) were used to recruit reads against pelagiphage genomes for assessing global abundances [82]. Coverage and Reads Per Kilbobase of each contig per Million Reads (RPKM) were calculated using coverm (available at https://github.com/wwood/CoverM) with thresholds on minimum read percent identity 90% and minimum covered fraction of 40% 'coverm contig –bam-files *.bam –min-read-percent-identity 0.9 –methods RPKM –min-covered-fraction 0.4'. To compare coverage threshold between HTVC010P and Skadi the coverm was used with minimum coverage thresholds set to 0 to 100%. The Ocean Data Viewer (ODV) v.5.2.0 available at (https://odv.awi.de/) was used to visualize read recruitment for Skadi. Median and mean RPKM values between different phage genomes were calculated within the R statistical software, using 95% confidence intervals (CI) and 1,000 bootstraps. Ordination analysis was performed using the coverm RPKM values and GOV2 metadata and the phyloseq [120] package for R v.4.0.3 (https://www.r-project.org/).

*Single Nucleotide Variant analysis of* Skadi *populations*

Metagenomic mapped reads to the Skadi genome were prefiltered to a mean coverage greater than 1 and detection greater than 70%. Single Nucleotide Variant (SNV) profiles with a minimum departure from base consensus value of 10%, and a minimum of 5X nucleotide coverage across the filtered metagenomes were generated using anvi-gen-variability-profile command from Anvi'o (version 7.1) package [121]. Dendrogram was generated based on a distance matrix representing pairwise comparisons as a function of SNVs. Dendrogram was visualized using the ggtree [122] package in R. The matching SNV profiles of the 461 variant positions were represented as a heatmap using ggplot package [123]. Figures were generated and edited with cowplot package (Wilke lab; https://doi.org/10.5281/zenodo.4411966) and inkscape image editing software (www.inkscape.org).

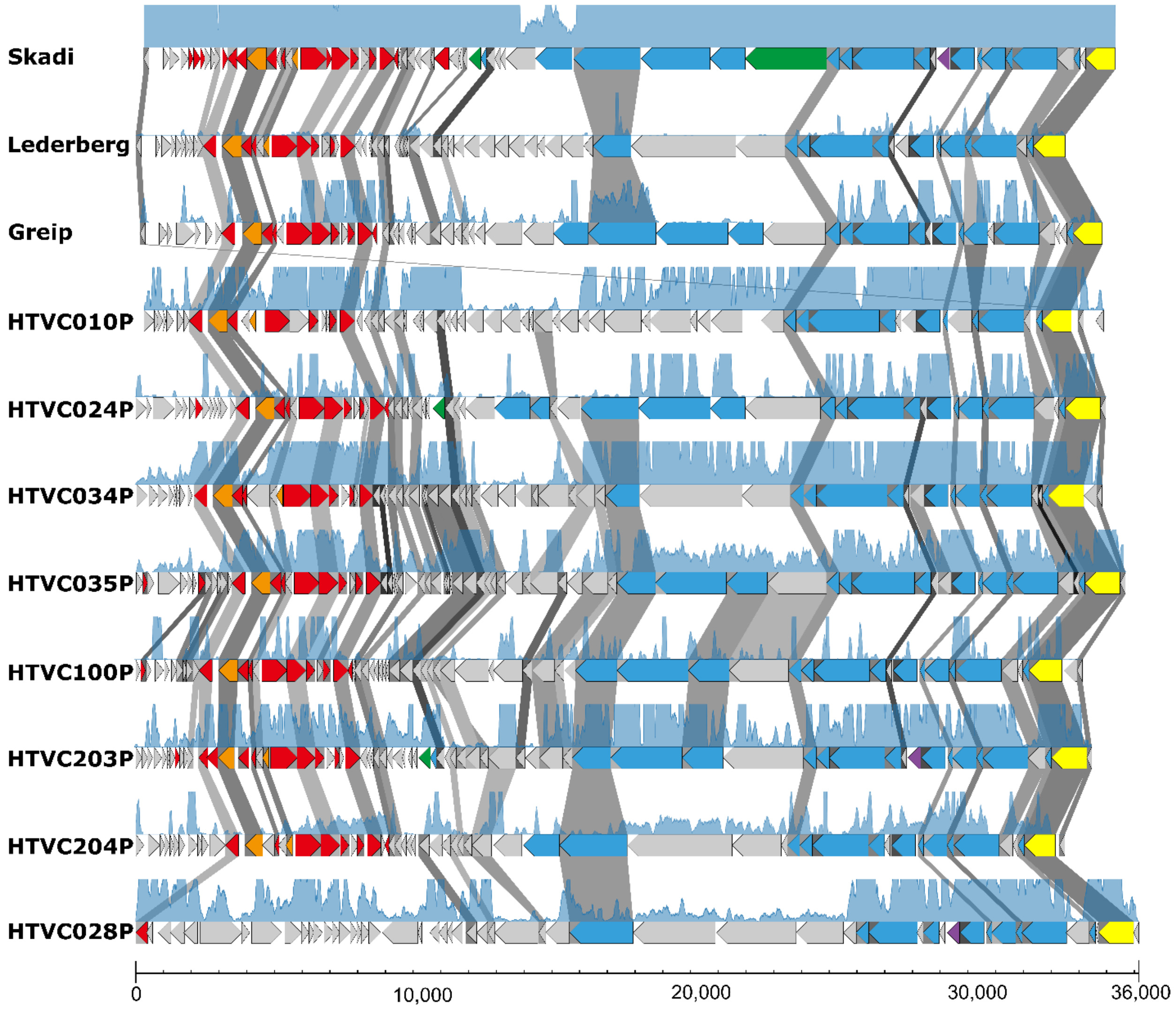


**Supplementary Figure 1. Genomic map of all members of the proposed *Ubiqueviridae* (HTVC010P-types and Skadi).** Arrow directions represents coding strand (positive/negative), shading connecting the open reading frames (ORFs) indicate identity between shared genes. The filled blue line graph indicates per nucleotide coverage (capped at 100) based on Global Ocean Virome (GOV2) reads mapped against phage genomes.

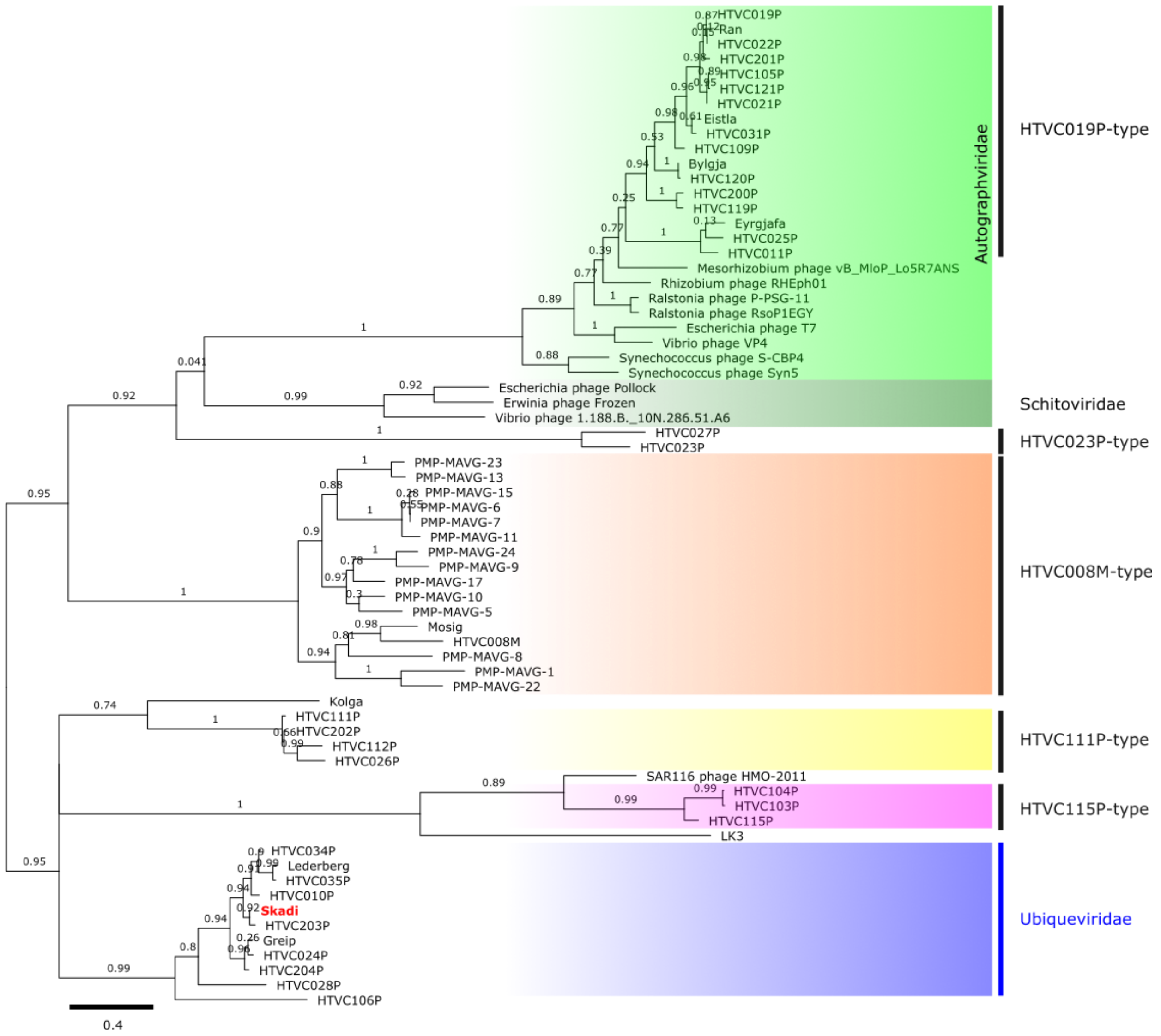


**Supplementary Figure 2 Phylogeny of pelagiphage *TerL* genes**. Unrooted neighbour-joining tree (100 bootstraps) of the *TerL* gene found in Pelagibacter phages and representatives of other viral families. Branches are coloured to highlight the different taxonomic groups in relation to the proposed *Ubiqueviridae*. Pelagibacter phage genome types are marked by black bars.

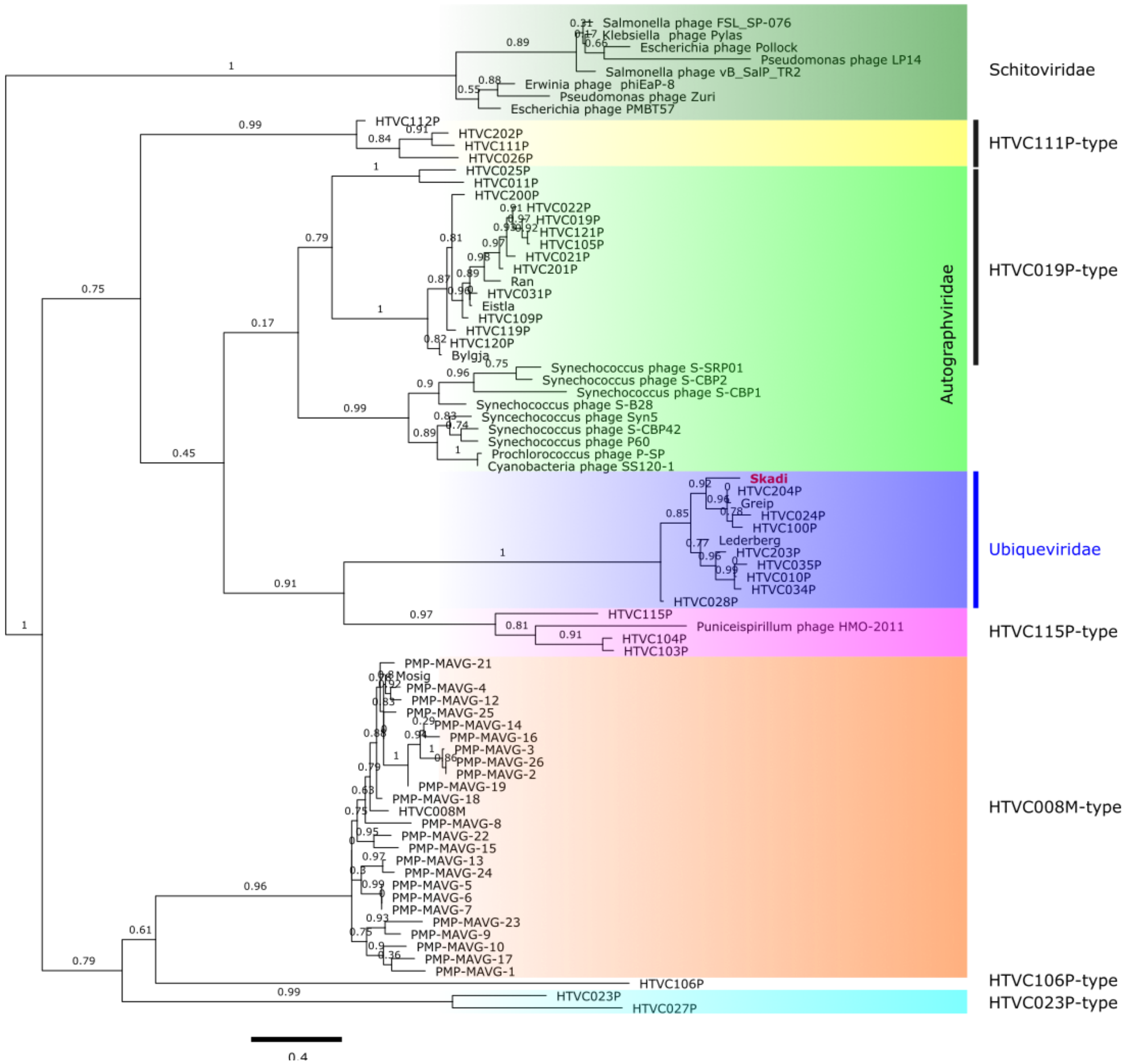


**Supplementary Figure 3. Phylogeny of pelagiphage tail tube B genes**. Unrooted neighbour-joining tree (100 bootstraps) of the tail genes related to Skadis tail tube B protein found in *Pelagibacter* phages and representatives of other viral families. Branches are coloured to highlight the different taxonomic groups in relation to the proposed *Ubiqueviridae*. *Pelagibacter* phage genome types are marked by black bars.

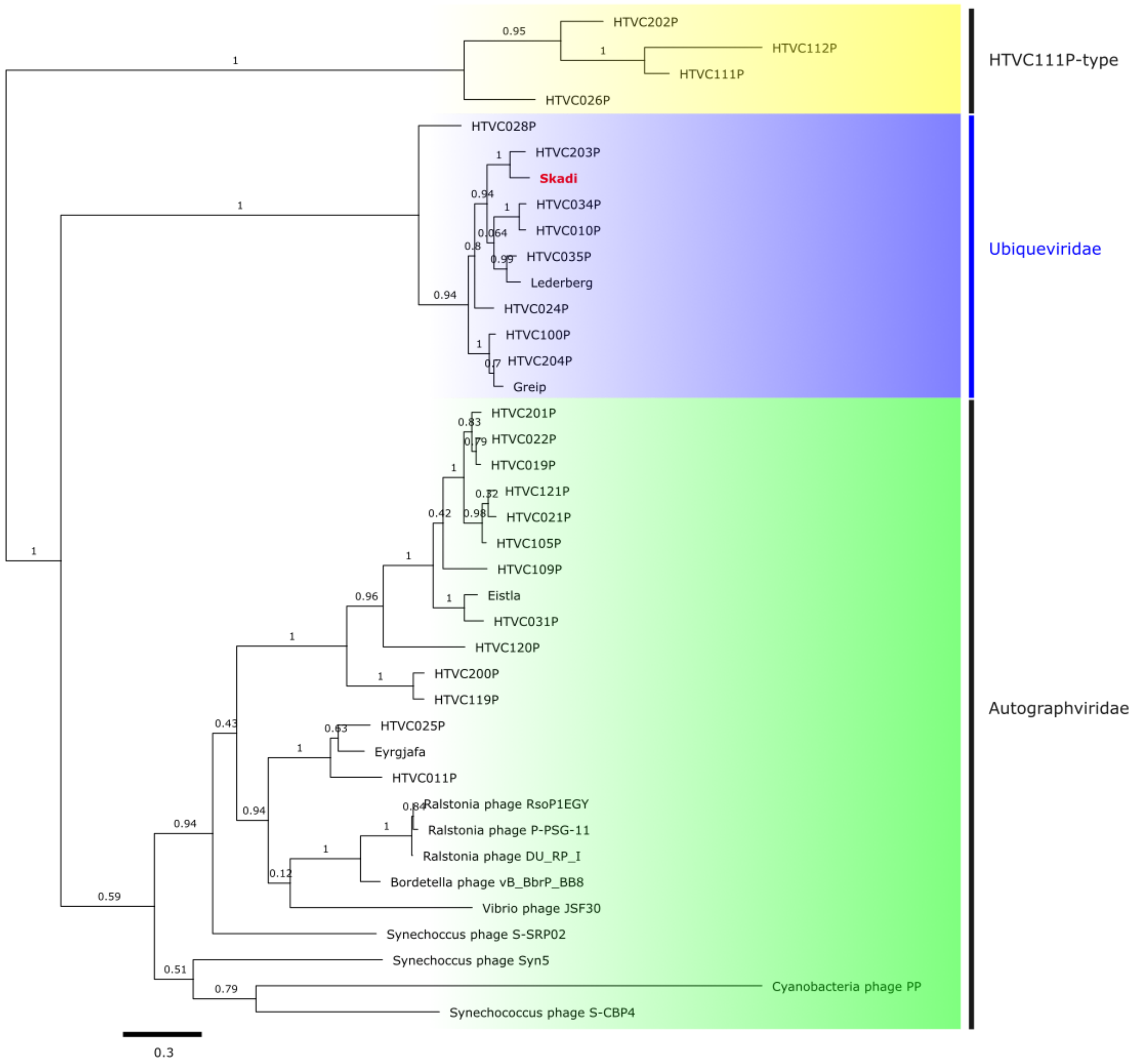


**Supplementary Figure 4. Phylogeny of pelagiphage major capsid protein genes.** Unrooted neighbour-joining tree (100 bootstraps) of the major capsid protein found in Skadi, *Pelagibacter* phages and representatives of other viral families. Branches are coloured to highlight the different taxonomic groups in relation to the proposed *Ubiqueviridae*. *Pelagibacter* phage genome types are marked by black bars.

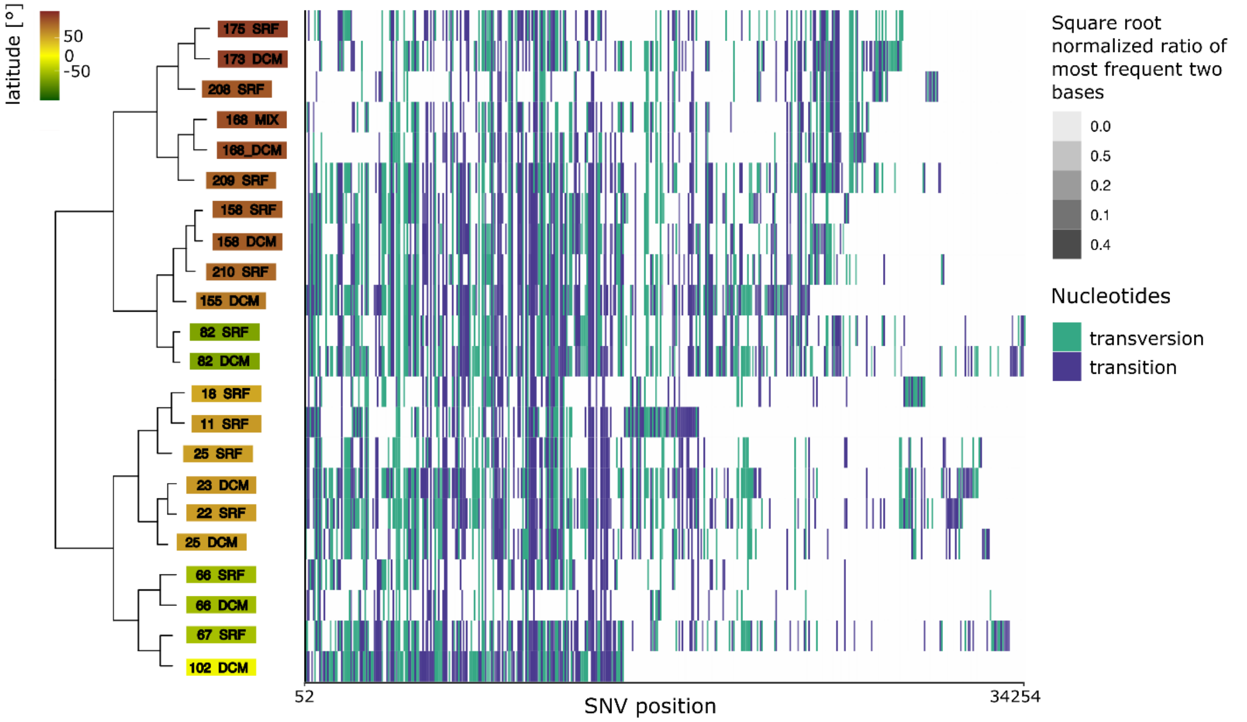


**Supplementary Figure 5. Dendrogram representing the distances between Single Nucleotide Variant profiles of the reads from the GOV2 metagenomes mapped to the Skadi genome.** A dendrogram representing the hierarchical clustering of the reads mapped to the Skadi genome is shown on the left side of the figure. Tips are color-coded based on the latitude where the sample was collected. A heatmap representing the profiles of the variable positions (461, columns) is shown on the right side matching the metagenome of the dendrogram tips. Nucleotide variants (compared to Skadi) are color-coded and shade of the cells represent the square-root normalized ratio of the two most frequent bases at the position (degree of variation in the position, lighter = more conserved- darker = more variable).

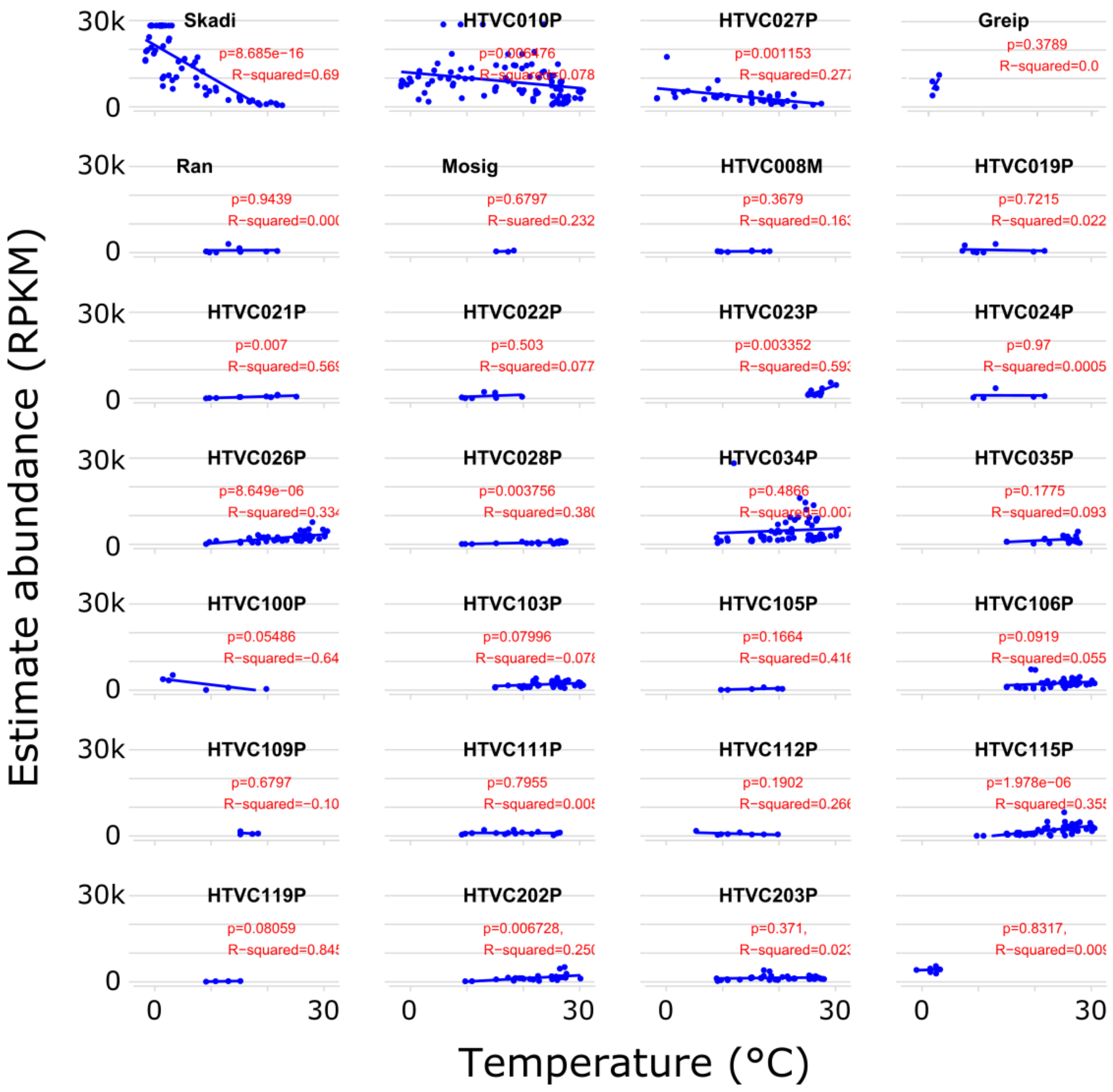


**Supplementary Figure 6. Pelagiphage Skadi correlates with Latitude**. Estimated abundances (RPKM) of isolated *Pelagibacter* phages from the GOV2 dataset with linear regression lines, samples with estimated abundances between zero and three RPKM in all viromes were excluded for clarity

**Supplementary Table 1.** Shared core genes identified in each genome of the *Ubiqueviridae*.

| | Gene ID number of phage strain: | | | | | | | | | | |
|---|---|---|---|---|---|---|---|---|---|---|---|
| Core Gene Annotation | Greip | HTVC010P | HTVC024P | HTVC028P | HTVC034P | HTVC035P | HTVC100P | HTVC203P | HTVC204P | Lederberg | Skadi |
| Tetratricopetide repeat | 8 | 15 | 48 | 37 | 50 | 55 | 40 | 54 | 45 | 10 | 50 |
| DNA binding | 9 | 14 | 49 | 38 | 51 | 56 | 51 | 55 | 46 | 11 | 51 |
| Acyltransferase | 10 | 13 | 50 | 40 | 52 | 57 | 52 | 56 | 47 | 12 | 52 |
| Tail tube B | 12 | 10 | 52 | 43 | 55 | 60 | 54 | 59 | 49 | 15 | 55 |
| Tail tube A | 13 | 9 | 53 | 44 | 56 | 61 | 55 | 60 | 50 | 16 | 56 |
| MCP | 14 | 8 | 54 | 45 | 57 | 62 | 56 | 61 | 51 | 17 | 57 |
| Tetratricopetide repeat | 15 | 7 | 55 | 46 | 58 | 63 | 57 | 62 | 52 | 18 | 58 |
| Portal | 16 | 6 | 56 | 47 | 59 | 64 | 58 | 63 | 53 | 19 | 59 |
| Hypothetical | 20 | 4 | 59 | 49 | 62 | 67 | 61 | 65 | 56 | 21 | 61 |
| TerL | 21 | 3 | 60 | 51 | 63 | 68 | 62 | 66 | 57 | 22 | 63 |

**Supplementary Table 2.** Overview of shared core genes per pelagiphage type.

| Taxon / Group | morphotype | core genes | Species |
|---|---|---|---|
| Ubiqueviridae | podophage | 10 | 11 |
| Autographviridae | podophage | 15 | 16 |
| Kolga-type | siphophage | n.a. | 1 |
| Mosig-type | myophage | 79 | 2 |
| HTVC111P-type | podophage | 28 | 4 |
| HTVC103P-type | podophage | 31 | 3 |
| HTVC023P-tpye | podophage | 33 | 2 |
| HTVC106P-type | podophage | n.a. | 1 |
